# Identification of divergent botulinum neurotoxin homologs in *Paeniclostridium ghonii*

**DOI:** 10.1101/2022.08.17.504336

**Authors:** Xin Wei, Travis Wentz, Briallen Lobb, Michael Mansfield, William Zhen, Huagang Tan, Zijing Wu, Sabine Pellett, Min Dong, Andrew C. Doxey

## Abstract

Botulinum neurotoxins (BoNTs) are the most potent family of toxins known to science. Bioinformatic studies in recent years have revealed that they are members of a broader toxin family, with an increasing number of divergent homologs identified in genomes of organisms outside of the *Clostridium* genus. Here, we report the identification of two putative divergent BoNT-like homologs in the genomes of two strains of *Paeniclostridium ghonii*. We designated them PG-toxin 1 (PGT1) and PG-toxin 2 (PGT2), which share ~54% protein sequence identity. Unlike any other known BoNT homologs, PGT1 and PGT2 are composed of two separate subunits encoded on two neighboring genes: one encoding the protease domain (light chain, LC) with a conserved HExxH motif, and the second encoding the heavy-chain (HC) containing the putative translocation domain and receptor-binding domain. Phylogenetic analysis of both the LC and HC reveal that it is a divergent member of the lineage of BoNT that also includes BoNT/X, BoNT/En and the insecticidal PMP1. The gene clusters harboring PGT1 and PGT2 also include a putative insecticidal delta-endotoxin, Cry8Ea1, as well as putative endolysin and bacteriocin genes that may facilitate lytic toxin secretion, suggesting a possibility that this gene cluster might serve an insecticidal purpose.

## Introduction

Clostridial neurotoxins (CNT) which include botulinum neurotoxin (BoNT) and tetanus neurotoxin (TeNT) are notable as the most potent toxins known to humankind, and they are the molecular agents responsible for botulism and tetanus neuroparalytic diseases. BoNTs target motor nerve terminals and enter neurons via receptor-mediated endocytosis, and then transfer their LC protease domains into the cytosol where they cleave host SNARE (soluble N-ethylmaleimide-sensitive factor attachment protein receptor) proteins including VAMP1/2/3, syntaxin 1, and SNAP25, resulting in blockage of neurotransmitter release [1–3]. TeNT shares the same mode of action with BoNTs, but it undergoes retrograde transport along the axon after entering motor neurons, reaching the spinal cord, getting released from the motor neuron soma, and then reentering inhibitory interneurons, eventually resulting in a distinct phenotype of spastic paralysis. Both forms of paralysis cause death due to respiratory failure.

BoNTs are typical A-B bacterial toxins composed of two polypeptide chains: the light chain (LC), which is the protease domain responsible for SNARE proteolysis, and the heavy chain (HC), which includes the translocation domain (Hn) that transports the LC across the endosomal membrane into the cytosol, and the receptor-binding domain (Hc) that binds to a number of different host cell receptors including polysialogangliosides, synaptotagmin I/II and glycosylated SV2, depending on BoNT serotype [3]. The LC and HC are encoded on a single gene and initially synthesized as a single polypeptide. The linker region between LC and HC needs to be proteolytically cleaved, by either bacterial proteases or host proteases, which is essential for the activity of the toxin. After the linker is cleaved, the LC and HC remain connected via a single interchain disulfide bond. This disulfide bond would be reduced once the LC reaches the cytosol, thus releasing the LC into the cytosol.

BoNTs were classified into seven serotypes (A-G), which include many subtypes [4, 5]. Serotypes A, B, E, and F are associated with human botulism, whereas subtypes C, D and hybrid CD are primarily associated with botulism outbreaks in wildlife. The host specificity of BoNT/G is less clear. BoNTs are produced not only by four groups of *C. botulinum* (Group I strains are human-associated and Group III are animal-associated), but also by strains of *C. noyvi, C. baratii, C. sporogenes*, and *C. butyricum*. BoNTs are encoded chromosomally, on plasmids, or on phages depending on the subtype and strain [6]. Related non-toxigenic strains are also common [7].

With the growing availability of genomes in public databases, researchers have identified a growing number of BoNT-related toxins bioinformatically (reviewed in [8–11]). In 2015, Mansfield et al. reported the first identification of a BoNT homolog outside of *Clostridium*, which occurred in the genome of the organism, *Weissella oryzae* [12]. This toxin, labeled “BoNT/Wo” was reported to cleave VAMP within a novel site [13]. Although BoNT/Wo conserves all BoNT domains, phylogenetically it sits on a divergent branch and outgroups the entire BoNT phylogenetic tree. This suggests that the lineage including BoNT/A-G may have evolved as a sublineage of a broader BoNT-like toxin family. Following the identification of BoNT/Wo, Zhang et al. identified BoNT/X in a *C. botulinum* strain, another distinct member of the BoNT family which formed its own lineage and serotype [14]. BoNT/X was demonstrated to cleave multiple VAMPs including several non-canonical substrates that cannot be cleaved by any other BoNTs. A year later, another BoNT-like toxin was discovered within a strain of commensal organism, *Enterococcus faecium* [15, 16]. This BoNT, labeled BoNT/En, was shown to cleave both SNAP25 and VAMP2 at unique sites, and reside on a common conjugative plasmid, suggesting a mode of circulation within *Enterococcus* strains. Interestingly, BoNT/En showed several similarities (phylogenetic and gene neighborhood structure) to BoNT/X, clustering with it in the BoNT phylogenetic tree. In 2019, a new family of toxins were also identified in the genome of *Chryseobacterium piperi*, which resembled BoNTs in terms of their domain structure [17]. These toxins have not been shown to cleave SNAREs and to date have unknown substrate specificity. However, their sequence and structural similarities to BoNTs suggest that they are more distant evolutionary relatives, even more so than the divergent BoNT homolog, BoNT/Wo.

Finally, the most recent addition to the BoNT family tree has come from studies of the insecticidal organism, *Paraclostridium bifermentans*. Contreras et al. demonstrated that this organism produces a BoNT-like neurotoxin, labeled PMP1, that has insecticidal activity toward anopheline mosquitoes by cleavage of the mosquito SNARE protein, syntaxin 1 [18]. PMP1 clustered within the divergent lineage of BoNTs including BoNT/X and BoNT/En, despite possessing only 34-36% identity to these toxins. In addition, PMP1 shares several genomic features with BoNT/X and BoNT/En, namely, the presence of neighboring *ORFX* genes, which were suggested to play a role in facilitation of PMP1 absorption in the insect gut, since their presence along with PMP1 increased its toxicity [18]. The identification of PMP1 as an insecticidal BoNT-like toxin aligns with earlier work which highlighted the numerous links between BoNTs and insecticidal pathogens [9]. Particularly, these links include the presence of homologs *ORFX* and *HA* genes in insecticidal gene clusters, the detection of partial homologs of BoNTs encoded within the genomes of entomopathogenic fungi, and the identification of BoNT-like gene fragments insect gut metagenomes [11].

In this work, we identified a novel BoNT homolog gene cluster in the organism, *Paeniclostridium ghonii*. The cluster is conserved in two distinct strains and also contains a neighboring gene encoding a putative insecticidal Cry toxin [19]. We analyzed these new BoNT homologs in terms of their sequence, structure, phylogenetic relationships to other BoNTs, and gene neighborhood. Our work adds to the growing diversity of BoNT-like toxins.

## Results and Discussion

### BLAST-based identification of BoNT-related genes in P. ghonii

On Jul 5, 2022, we performed a BLAST-based search for clostridial neurotoxin homologs in the NCBI database, restricting our search to organisms outside of the *Clostridium* genus. Our search, using BoNT/A1 (accession # WP_021136346.1) as a query identified two previously unidentified matches in the genome of organism, *Paeniclostridium ghonii* (strain NCTR 3900). The BoNT/A1 light chain (LC) region encoding the metalloprotease domain matched a protein labeled as “M91 family zinc metallopeptidase” (WP_250673407.1) with an *E*-value of 1e-21 and 34% amino acid identity. The BoNT/A1 heavy chain (HC) region matched a second protein labeled as “hypothetical protein” (WP_250673408.1) with an *E*-value of 1e-54 and 28% amino acid identity. We designated these proteins “PG-toxin1-LC” and “PG-toxin1-HC” or the short forms, “PGT1-LC” and “PGT1-HC”.

Next, we examined the JGI genome database for additional strains of *P. ghonii*, and found one additional genome (unique from NCTR 3900), labeled as strain “DSM 15049”. A similar toxin gene cluster was identified in the genome of this strain, including orthologs of both proteins identified in NCTR 3900. Interestingly, these orthologs possess 39% identity for the LC protease gene and 61% identity for the HC gene, suggesting possible rapid divergence of the toxin sequences since its acquisition by an ancestral *P. ghonii* strain similar to those observed for BoNTs in Group I and II *C. botulinum*. We designated these proteins “PG-toxin2-LC” and “PG-toxin2-HC” or the short forms, “PGT2-LC” and “PGT2-HC”.

A table of pairwise percentage identities to a representative subset of BoNT-like proteins is shown in **Figure 1a** for the LC and **Figure 1b** for the HC. Overall, combining their LC and HC sequences, PG-toxin1 and Pg-toxin2 share ~54% sequence identity. The PG-toxin LC and HC sequences consistently show the highest percentage identities to PMP1, BoNT/En and BoNT/X compared to other BoNT-like sequences, suggesting a possible phylogenetic affiliation with this lineage.

**Figure 1.**
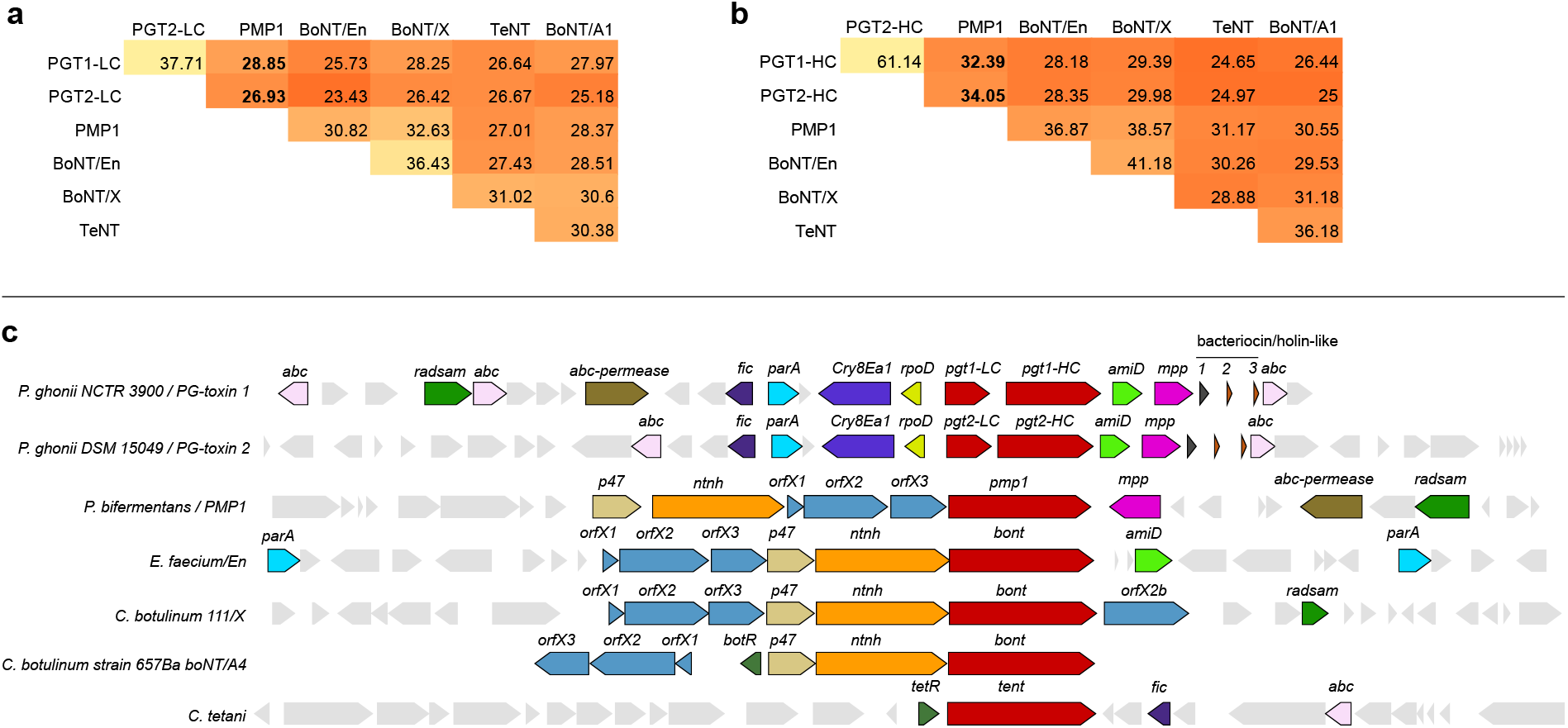
Identification of BoNT-like genes in genomes of two *Paeniclostridium ghonii* strains. Pairwise amino acid sequence identities between PG-toxin-LC (a) and PG-toxin-HC (b) and a subset of BoNT proteins. (c) Genomic neighborhoods surrounding PG-toxin genes and other BoNT genes.

### Gene neighborhood analysis

Next, we investigated the genomic neighborhoods surrounding both PG-toxin genes (**Figure 1c**). In both *P. ghonii* genomes, the PG-toxin-LC gene encoding the predicted zinc metalloprotease is immediately upstream of the PG-toxin-HC gene encoding the translocase and binding domain. To verify that this is not due to sequencing errors, we analyzed the region further and determined that the intervening region is non-coding with numerous stop-codons in all reading frames.

To examine the source of the fragment (chromosomal versus plasmid), we analyzed the gene content up and downstream of the PG-toxin gene cluster. Within the scaffold from DSM 15049 containing the PG-toxin2 genes, which was more complete (64,948 bp) than the toxin-encoding contig from NCTR 3900 (29,173 bp), we identified genes encoding plasmid replication initiation proteins. BLAST analysis of these proteins confirmed a close relationship to orthologous genes from *P. bifermentans*, suggesting a link to the toxin-encoding megaplasmids of *Paraclostridium* [18]. To investigate the source of the contigs in their entirety, contig level assemblies were passed through rfPlasmid (v0.0.16) using a *Clostridium* model and default settings. The contigs carrying the BoNT-like gene sequences for *Paeniclostridium ghonii* DSM15049 and NCTR 3900 were predicted as plasmids with 73.28% and 74.66% percent of the vote, in excess of the 40-60% threshold where incorrect classifications typically fall (rfPlasmid). Similarly, another near-*Clostridium* species with a known BoNT-like sequence on a reported circularized plasmid, *Paraclostridium bifermentans* parabai, was predicted as a plasmid with 79.16% of the vote. We therefore speculate that the toxin loci may reside on a plasmid.

The PG-toxin1 locus from strain NCTR 3900 is similar but non-identical from that of the PG-toxin2 locus from strain DSM 15049. A notable unique feature of the PG-toxin locus is the presence of a nearby gene encoding an insecticidal Cry delta endotoxin (accession # WP_250673405.1). A cry toxin prediction was confirmed by the Conserved Domain Database, with matches to all three domains (N, M, and C-terminal region) with significant *E*-values all less than 1e-15. BLAST analysis of this protein also confirmed its relationship to a Cry8Ea1 family insecticidal delta endotoxin from *Bacillus thuringiensis* (accession # WP_172452114.1) with ~40% amino acid identity. The closest sequence of known structure is *B. thuringiensis* cry3Aa (PDB ID 4QX0_A) at 32% identity. In both P. ghonii genomes, this Cry toxin gene is encoded in the opposite direction to the PGT-LC genes, which suggests that it may be regulated differently.

In addition, The PG-toxin locus also encodes several genes shared with those found in other BoNT gene clusters. These include *amiD* (n-acetylmuramoyl-l-alanine amidase gene) and *parA* genes which are also present in the BoNT/En locus of *E. faecium;* and *abc* (ABC transporter ATP-binding protein) and *fic* (filamentation induced by cAMP protein) genes that are also found in the *tent* locus of *C. tetani*. Three nearby genes are also shared with the PMP1 locus of *P. bifermentans:* an *mpp* gene encoding a putative metallophosphatase family protein, an *abc-permease* (ABC transporter ATP-binding protein/permease) and a *radsam* gene encoding a radical SAM domain containing protein. Interestingly, the *orfx, ha* and *ntnh* genes present in other BoNT gene clusters are absent. Downstream of the *mpp* gene in both *P. ghonii* genomes is a cluster of genes encoding a BhlA/UviB family holin-like peptide (labeled “1”) and two genes encoding aureocin-like type II bacteriocins (labeled “2” and “3”) (**Figure 1c**). Other toxins are known to be exported via holin-dependent mechanisms [20].

### Protein sequence and domain analysis

Next, we compared the PG-toxin sequences to other BoNTs to examine the presence/absence patterns of key motifs and structural features important for neurotoxin activity. While typical BoNTs are proteolytically cleaved into two distinct chains (LC and HC) that remain connected via a disulfide bond (**Figure 2a**), a unique feature of the *P. ghonii* BoNT-like toxins is the split of the LC and HC into two distinct coding sequences (**Figure 2b**). The correspondence of the PG-toxin-LC and neighboring PG-toxin-HC sequences with the LC and HC of BoNTs can also be seen at a structural level, as revealed by the homology models shown in **Figure 2b**.

**Figure 2.**
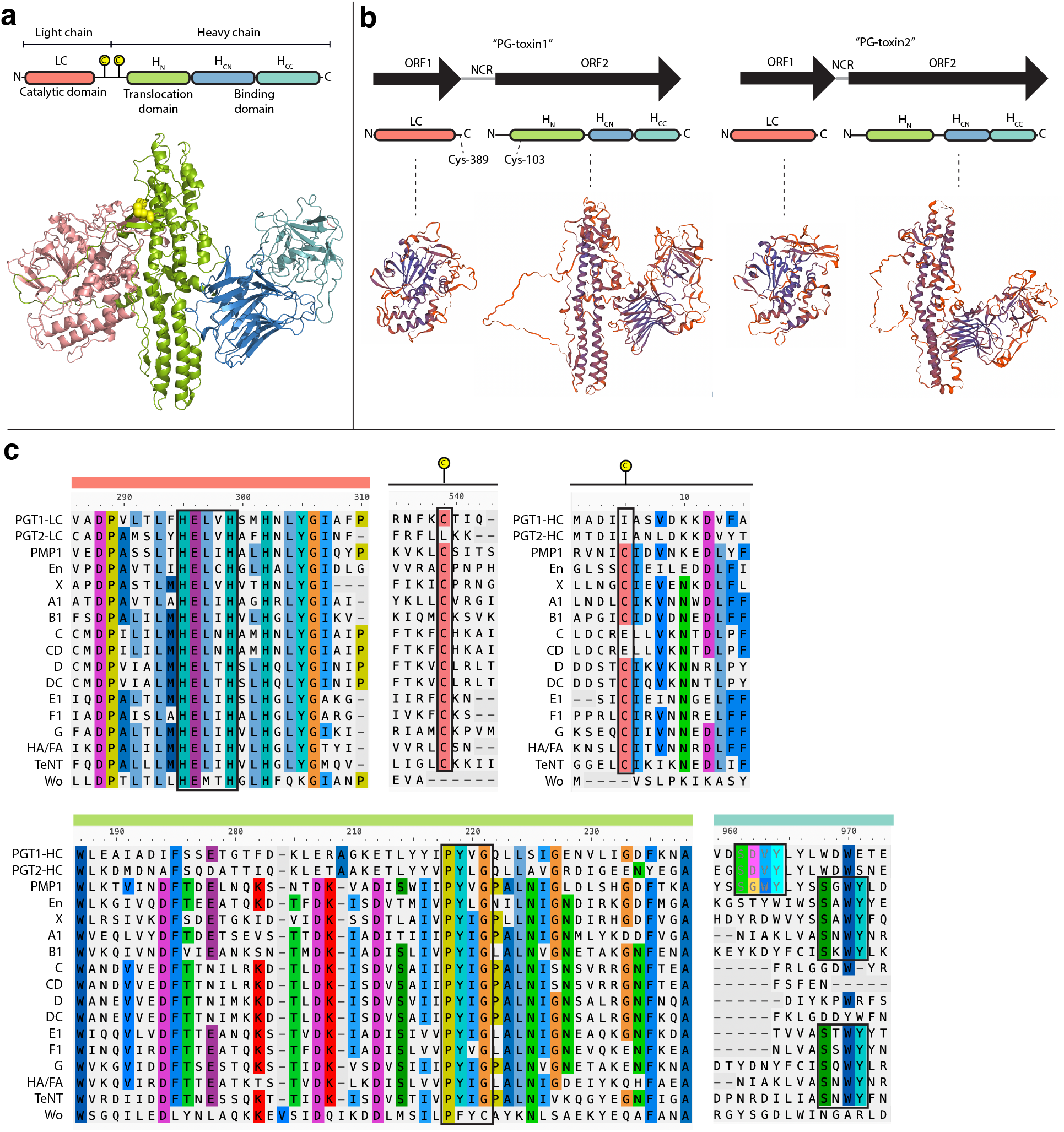
Structural and motif analysis of PG-toxins in relation to BoNT. (a) Domain structure of BoNT/A1 (above) depicting the light and heavy chain and corresponding crystal structure (below) from PDB 3BTA. (b) Protein domain architectures of PG-toxin open reading frames (ORFs). *P. ghonii* strain NCTR 3900 and DSM 15049 both encode two ORFs separated by a non-coding region (NCR) which correspond to the LC and HC regions of BoNT, which are collectively referred to as PG-toxin1 and PG-toxin2, respectively. Structural models of these regions confirm their correspondence to the BoNT LC and HC. (c) Selected amino acid alignments depicting key functional residues of known importance in BoNT. The catalytic HExxH motif required for zinc endopeptidase activity is conserved in PGT1-LC and PGT2-LC, while all but one of the canonical Cys residues involved in the LC-HC disulfide bond are missing. The PxxG motif within the translocation domain is also conserved, and the canonical SxWY ganglioside binding motif is absent, although a second upstream SxxY motif is present in an equivalent site to a second SGWY motif previously identified in PMP1.

The gene split between the LC and HC raises the question of how these domains may physically interact if they are assumed to function equivalently to BoNTs. Interestingly, PG-toxin1-LC contains a Cys residue at its C-terminal region (Cys-389), whereas this Cys residues appear to be absent in PG-toxin2-LC. The closest second Cys residue in PG-toxin-1-HC occurs at residue Cys-103 but this is within the translocase domain. Thus, it appears unlikely that the PG-toxin-LC and HC domains interact via the same disulfide bond as in BoNTs. Structural models of the PG-toxin chains (**Figure 2b**), however, may provide some insights into a possible interaction between the LC and HC domains. That is, the “belt” region of the translocase domain in the HC domains, which is known to wrap around the LC in other BoNTs [21, 22], is present in both PG-toxin1-HC and Pg-toxin2-HC. We therefore speculate that this belt may provide a mechanism for forming a complex between the light and heavy chains, although this hypothesis remains to be evaluated experimentally.

Both LC protease sequences (PG-toxin1-LC and PG-toxin2-LC) contain the conserved HExxH motif required for zinc-endopeptidase activity, and more specifically an HELVH motif identical to that in BoNT/X (**Figure 2c**). Overall, PG-toxin1-LC and PG-toxin2-LC regions possess a maximum of 28.9% and 26.9% identity to their closest BoNT-like homolog (PMP1), respectively (**Figure 1a**).

A region of sequence conservation for the translocation domain is also shown in **Figure 2c**. The PxxG motif, which has previously been identified as a strongly conserved motif across BoNTs (including divergent homologs) [17] is conserved in the PG-toxin translocation domains. This PxxG motif is also conserved in translocation domains of diphtheria toxins (DT) and DT-like homologs [23], as well as large clostridial toxins (LCTs) [24]. Overall, the putative translocation domains from PG-toxin1-HC and PG-toxin2-HC share ~48.2% identity, and possess a maximum of ~30% to other BoNT-related proteins (BoNT/En).

Lastly, we examined the C-terminal binding domain (H_cc_) of both PG-toxin HC sequences. In the H_cc_ domains of BoNT/A, B, E, F, G, X, En, PMP1, and TeNT, a conserved SxWY gangloside binding motif (which we designated “motif 1”) has been identified that is known to bind ganglioside cell receptors [25]. Importantly, in the mosquitocidal PMP1, a second SxWY motif (“motif 2”) is present immediately upstream of the canonical SxWY motif, as well as a third SWYG motif (“motif 3”) which was found to be critical for toxicity [18]. Interestingly, both PG-toxin-HC sequences lack the canonical SxWY motif (“motif 1”) but possess an SDVY motif in place of motif 2, which we hypothesize may play a role in receptor binding (**Figure 2c**).

### Phylogenetic analysis

To explore their evolutionary relationships to other BoNT proteins, we separately aligned the PG-toxin LC and HC protein sequences to corresponding regions from a representative set of available BoNTs retrieved from https://bontbase.org (also including PMP1). We then built maximum-likelihood phylogenies for the LC (**Figure 3a**) and HC (**Figure 3b**).

**Figure 3.**
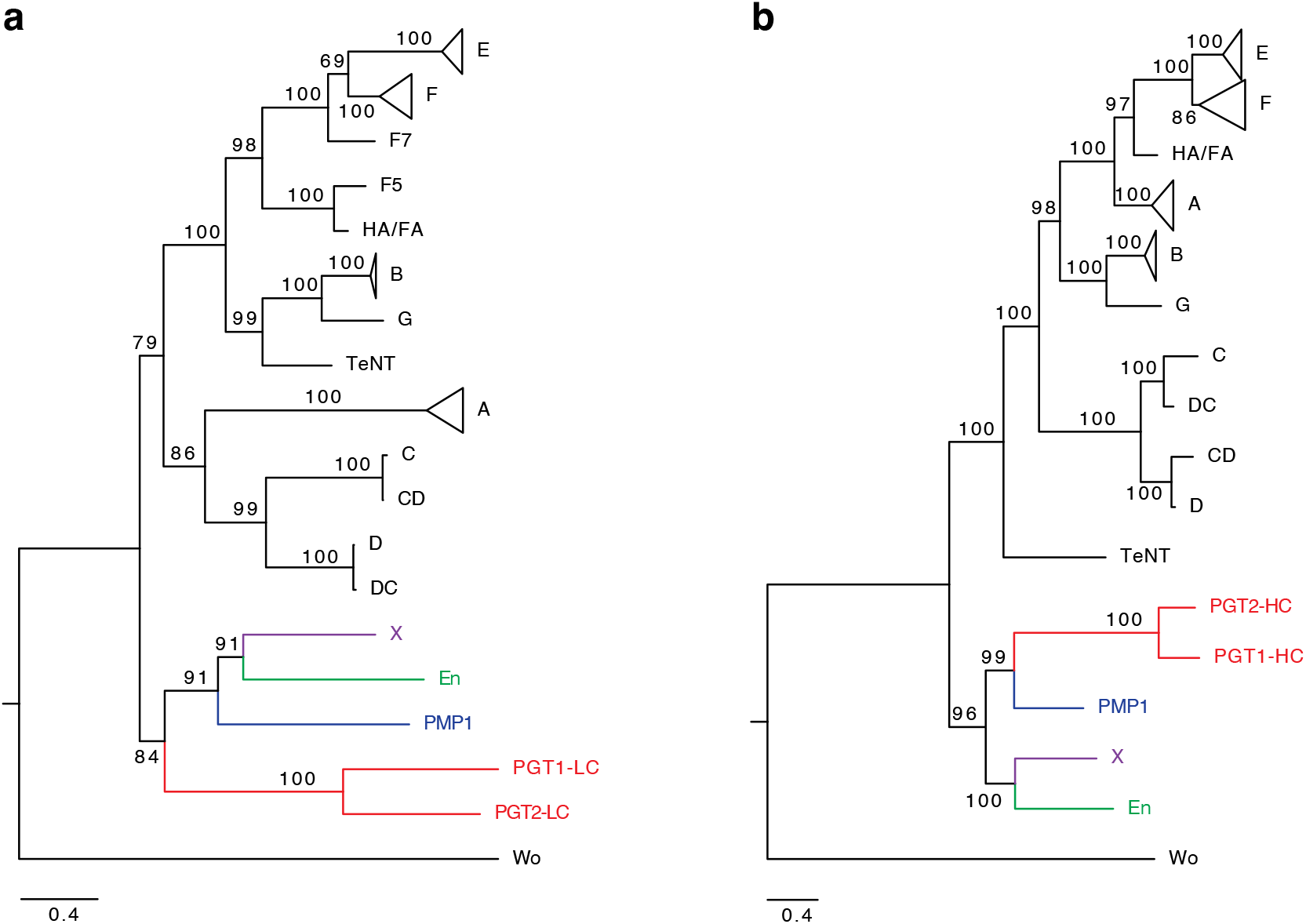
Phylogeny of the BoNT family including PG-toxin. (a) Phylogeny of the LC encoding the protease domain. (b) Phylogeny of the HC encoding the translocation domain and binding domain. In both trees, PG-toxin-LC and PG-toxin-HC cluster with the X/En/PMP1 lineage. Bootstrap values are indicated for all major clades.

The phylogenetic placement of the PG-toxin-LC and PG-toxin-HC sequences was similar. In both cases, they clustered with the X/En/PMP1 lineage (**Figure 3**). The PG-toxin-LC clustered as a sister lineage to X/En/PMP1 with moderate (84%) clade support, whereas the PG-toxin-HC clustered specifically with PMP1 with very strong (99%) clade support. As expected, the PG-toxin genes from both *P. ghonii* strains clustered as neighbors indicating a common origin in an ancestral *P. ghonii* strain.

### Summary and phylogenetic analysis of the organism, Paeniclostridium ghonii

*Paeniclostridium ghonii* was first identified in the 1930s and was originally named *Clostridium ghoni* (Prévot, 1938). It was later reclassified as a member of the genus *Paeniclostridium* [26]. The organism has been found in soil, marine sediments, rumen, soft-tissue infections in humans and human faeces, and in alfalfa silage [27]. The species is also notable as it was identified as a member of the microbiome in the colon of the Iceman [28]. One of the most closely related species is *Paeniclostridium sordellii* (*Clostridium sordellii*), a Gram-positive pathogen that causes severe oedemic, myonecrotic or enterotoxic infections in humans, and farm animals [29].

To examine the phylogenomic similarities of *P. ghonii* to other available sequenced organisms, we retrieved genomes of all available *Paeniclostridium* species, as well as [Eubacterium] tenue and members of the genus, *Paraclostridium*, given their previously identified relationships to *P. ghonii* based on 16S analysis [26]. Based on genome-wide average nucleotide identity (ANI) clustering, the two strains of *P. ghonii* cluster as a sister lineage to a clade of *Paraclostridium* which includes *Paraclostridium bifermentans* and *P. benzoelyticum* (**Figure 4**). *P. ghonii* displays ANI values of ~87.4-87.6% to these *Paraclostridium* species and lower ANI values to [Eubacterium] tenue (83.4%) and *Paeniclostridium sordelli* (82.7%). The relatively close relationship between *P. ghonii* and *P. bifermentans* is noteworthy given the presence of BoNT-like genes associated with these species, and suggests the existence of a broader lineage of potential *Paraclostridium* and *Paeniclostridium* species that encode BoNT-like toxins.

**Figure 4.**
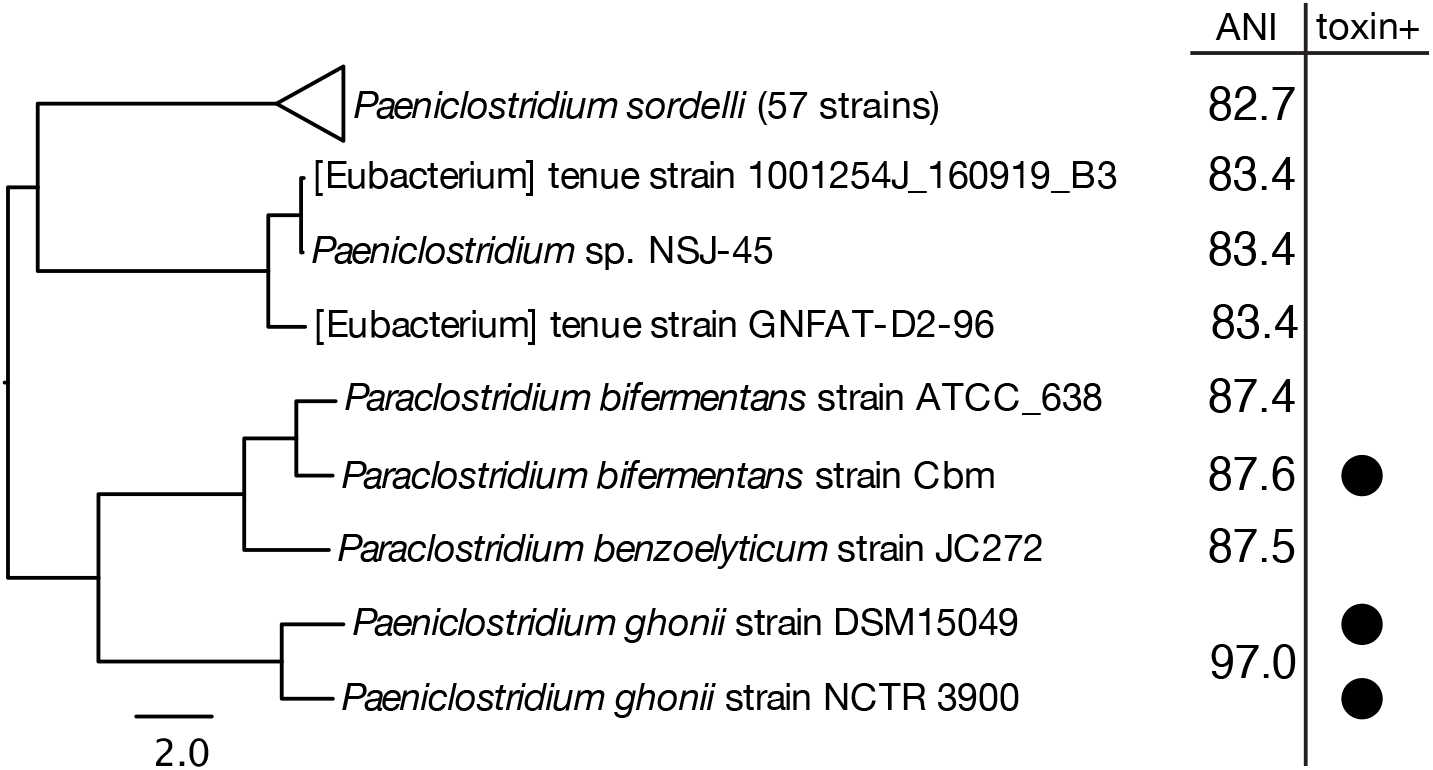
Phylogenetic clustering of *P. ghonii* genomes with the top 64 closest genomes in the NCBI database based on average nucleotide identity. Average nucleotide identity (ANI) values are shown on the right of the tree, and are averaged for the two *P. ghonii* strains.

## Methods

### PG-toxin identification

On Jul 5, 2022, the BLAST webserver (https://blast.ncbi.nlm.nih.gov/Blast.cgi?PAGE=Proteins) was used to perform a search for clostridial neurotoxin homologs in the NCBI nr protein database, restricting our search to organisms outside of the *Clostridium* genus. We used BoNT/A1 (accession # WP_021136346.1) as a query with default parameters.

### Phylogenetic analysis of PG-toxins

An alignment was made using sequences from bontbase.org as well as PMP1 (WP_150887772.1) using MAFFT with the L-INS-i algorithm [30]. The alignment was then split into the heavy and light chain regions corresponding to domain boundary definitions from PDB 3BTA. Conserved alignment blocks were then manually selected and used to produce two separate phylogenetic trees with the IQ-TREE algorithm [31].

### Structural modeling

The PG-toxin LC and HC sequences were submitted to the SwissModel resource for automated protein homology modeling [32]. Structural models were generated using a variety of template structures of existing BoNT related proteins. For PGT1-LC, PDB ID 6bvd.1.A (Botulinum Neurotoxin Serotype HA Light Chain) was used as the top identified template, which had 28.09% identity. For PGT2-LC, PDB ID 5bqn.1.A (Botulinum neurotoxin type D) was used as the top identified template, which had 26.36% identity. For PGT1-HC, 6g5g.1.B (Botulinum neurotoxin type B) was used as the top identified template, which had 25.32% identity. For PGT2-HC, 7by5.1.A (tetanus toxin) was used as the top identified template, which had 26.28% identity. Structural models were colored by QMEANDisCo Local quality estimate scores.

### Phylogenetic analysis of P. ghonii based on average nucleotide identity

We chose the closely related species of *Paeniclostridium ghonii* to build the species phylogeny, including all species in the *Paeniclostridium* genus, as well as *Paraclostridium bifermentans*, which was shown to contain PMP1 toxin, and *Paraclostridium benzoelyticum* that was reported as the species with the highest ANI in the NCBI database. The assembly files in .fna format of all the genomes were download from the NCBI (https://www.ncbi.nlm.nih.gov/assembly) and JGI database [33]. Then, the fastANI program was used to compute pairwise ANI and generate a distance matrix with identity values for the list of the assemblies [34]. The R package phangorn was used to build a phylogenetic tree using the neighbor-joining method based on the ANI distance matrix [35]. The ANIs for each of closely related genome to both *P. ghonii* genomes were averaged and indicated in **Figure 4**.

### Genomic neighborhood analysis

The genomic neighborhoods for ±20kb surrounding the neurotoxin genes were obtained from the NCBI database in .gff format. For PG-toxin genes, we downloaded the complete scaffold to collect as much sequence information as possible. The .gff files were uploaded to an in-house tool called AnnoView for gene neighborhood visualization (manuscript in preparation).

## Notes

### Competing Interest Statement

The authors have declared no competing interest.

